# Using Biometric Data to Measure and Predict Emotional Engagement of Video Games

**DOI:** 10.1101/2022.02.28.482337

**Authors:** Janette Vazquez, Samir Abdelrahman, Chris Wasden, Stuart Jardine, Colby Judd, Matthew Davis, Julio C. Facelli

## Abstract

Measurement and prediction of engagement in video games are very important because these are indicators to guide the development of game applications. Existing methods to predict game engagement are mostly based on subjective evaluations. This paper presents results correlating subjective and biometric data to show the potential of using biometric measures for assessment and prediction of engagement. Using three different games, we measured the biometric responses of participants and analyzed the responses using statistical and data mining methods. We compared the results with those obtained using subjective evaluations. Our results show that biometric measurements do correlate with subjective measures and better predict the self-reported engagement of a game.

## 1. Introduction

Gamification is a recent trend that applies game approaches to non-game fields to engage audiences in applications that will provide benefits and be fun to engage in. Fields such as health, business, marketing, and e-learning have taken advantage of gamification. For instance, there is a growing interest in the “gamification” of health care services leading to capturing more engagement from patients to enhance compliance with pharmaceutical and behavioral interventions. For instance, gamification’s overall goal in health care is to deliver interventions to an individual in new ways that keep individuals in treatment regimens that may otherwise be ‘demotivating’ or hard to maintain. However, bringing games to market requires substantial investments; therefore, return on investment can be greatly improved through developing methods to predict which games will be more engaging (1) by testing them early on in relatively small cohorts.

Numerous papers have reported on the evaluation of the effectiveness of health care games as well as the limitations and challenges currently facing the gamification of e-Health applications among others. There are two critical factors for the success of a game application, one is adoption, and the second is effectiveness. For instance, in a systematic literature review, various combinations of game elements utilized in e-Health were identified, as were the potential benefits and challenges that surround their integration into effective interventions. The challenges most frequently mentioned were user engagement and motivation, which need to be sustained in the long term for a game to be able to provide the benefits and therapeutic effects a gamified health application promises to deliver (2).

Current methods of evaluating games are through both subjective and objective measures, with the most common methods of measuring engagement for design decisions being through subjective self-reports, such as questionnaires and interviews, and objective reports, such as observational video analysis (3). While these objective methods appear to be preferable for being less biased, they are very labor-intensive, requiring extensive annotations of the videos by specialists. Coding gestures, body language, verbal comments, and other data as an indicator of human experience is a lengthy and rigorous process that has to be undertaken with great care, with researchers acknowledging biases, addressing interrater reliability, and not reading inferences that are not present (3). Subjective methods like the Self-Assessment Manikan (SAM) (4) are less labor-intensive but they are subject to significant biases (5). Subjective measures, although generalizable, convenient, and easy to gather tend to not find complex patterns, and participant’s responses may not correspond to the experience an individual had, introducing undesirable biases in the studies. Moreover, knowing that answers are being recorded, answers may sometimes be what participants believe the administrator wants or wants not to hear (3).

With the development of better and less costly biometric measurement technology, it is now possible to routinely collect biometric data feedback that could be sued to analyze gameplay and engagement without the need for trained researchers to commit a large amount of time into coding observational data from video recordings. Previous studies have used biometric data to show how physiological measures correlate with surveys and player experience (6-8). For example, Mandryk et al. and Yannakakis et al. have examined the correlation between several physiological measures (GSSR, EMG, respiratory and cardiovascular measures) with player’s experiences in computer games (3, 9). Earlier studies also focus on transforming GSR, HR, and other biometric data into arousal and valence using fuzzy models (10). Recent studies have used biometrics to measure what motivates a player in a game (11) or have focused on finding ways to change the playthrough of a video game to adapt to a person’s physiological responses, better known as affective gaming (12). Studies have also used biometric feedback for game user research (GUR) to see if user experience issues could be found during a playthrough that could then be used to improve the game design (13, 14). Another study focused on the different emotional responses created when playing four very different games and found that they resulted in different measures of valence, arousal, and discrete emotions for each game (4). Different levels of experience of test subjects have also been considered in studies (6). All these studies used biometric measures to assess different aspects of video games. None of these studies, however, formed a quantitative model for biometric user testing or analyzed whether biometric feedback of a finished video game could be used to assess the engagement level an individual reported to feel from a game.

In this study, we perform two analyses. In our first analysis, we look at the physiological measures and find if they correlate (or correspond) to subjective reports, similar to the approach followed by Mandryk et al. (3, 10). In our second analysis, we test the hypothesis that biometric data acquired during gameplay is a better determinant of engagement than traditional user survey-based testing measures, such as the GEQ. We collected three different sets of biometric measures: heart rate from the electrocardiogram, a GSR measure, and facial expression analysis. The biometric and GEQ datasets were analyzed using three different, but well-established, machine learning analysis methods (Support Vector Machine, Bayesian Networks, and Random Forests) to find which one of these has better predictive power. This work is an attempt to develop a well-established protocol for data collection and analysis toward the establishment of reliable quantitative methods in the prediction of engagement. To our knowledge, no previous research uses physiology as an indicator of fun or engagement with entertainment technology.

## 2. Methods

Participants were recruited and selected from a university and its related community with the approval of the research protocol by the University of Utah Institutional Review Board. Surveys were used to gather demographic data. A total of 40 participants were recruited. There were 47% female and 53% male, with a mean age of 35 years, which is representative of game users (1). Participants were then classified as falling into one of three levels of video game proficiency based upon experience: inexperienced gamers (< 2 hours of gameplay a week), casual gamers (2 – 10 hours of gameplay a week), and experienced gamers (> 10 hours of gameplay a week). These distinctions were made based on previous studies that found a difference in emotional engagement of experienced gamers versus casual players when using biometric measures (3, 15). Of the 40 participants selected, data on 12 participants were excluded from the analysis due to errors in the hardware in delivering consistent and reliable biometric signals to measure and analyze, resulting in 28 subjects for the final analysis. These 28 participants had a mean age of 21.7 ± 3.6, consisted of 60.7% males and 39.3% females, from which there were 11 casual gamers, 5 non-gamers, and 12 experienced gamers. Two of the subjects had experience playing two of the three games used in the study (Doom and BattleBorn), three others were familiar with Doom, and two others were familiar with Dead Effect 2.

Three games were selected for this study (supplementary material). After an analysis of previous studies on game selection, it was decided that a first-player shooter game genre would be most appropriate for this study due to its highly immersive nature. The study also chose not to use the multi-player version of these games to remove the engagement dynamics associated with this additional variable. By using only one genre, the study was able to eliminate genre as a variable that could create heterogeneity in the data, albeit perhaps losing some degree of generalizability. All three games were published and commercially released at about the same time and available on Steam, a large digital gaming platform, that also provided measures of game downloads, release date, and the number of plays within the last two weeks (16). The three games selected for the study had different levels of downloads and gameplay statistics.

As of June 8, 2021, Dead Effect 2 had 100,000 owners, with 29.6% of players engaging in the game longer than 5 minutes and an average playtime of 6:54 minutes. Doom had an average of 3,767,000 owners with an average playtime of 14:09 minutes with 64.79% of players engaging in the game longer than 5 minutes. BattleBorn had 935,000 owners with an average playtime of 10:02 minutes and 33.3% of players engaging in the game for 5 minutes or longer (16). Based on the most recent results of engagement and game playtime, we have categorized Doom as the most engaging game of the three, followed by BattleBorn, and then Dead Effect 2.

The Shimmer GSR+ device was used to collect data on heart rate and skin conductance (17). The device has an operating range from 22 kOhms to 680 kOhms with a standard error of 3% across that range. The iMotions (18, 19) software was used to collect all the data from facial expression analysis and was synchronized with the shimmer device (20). The GSR+ Shimmer device used the Ag/AgCl electrode positions against the palmar region of the medial and distal phalanges of the right hand, which was also the same hand that subjects used for the mouse that controlled some player features and actions of the game. The left hand of the subject was used to control keys on the computer keyboard when playing the game. The temperature of the study room was maintained at the same level throughout the study to ensure that the device was not affected by changes in room temperature that could cause variations in sweat gland production.

The Affectiva facial recognition software was used in conjunction with the iMotions software for facial expression analysis (21).

Participants were tested for the study on an individual basis during sessions that were approximately one hour long. The participants provided consent for the study upon arrival and were then given a detailed explanation of the experiment that they were about to participate in. After signing the consent forms, the participants completed a survey that collected basic demographic data, along with their experience playing video games and their experience with these three games. The participants were then guided to the study area, which was an empty office space that only included the equipment that was being used for the study. The participants were fitted with the Shimmer GSR+ device and were set up for the study to ensure that their facial expressions could be reliably recorded. The lighting in the room was maintained throughout the duration of the study and the room was evenly lit so participants experienced constant conditions.

The desk for the study supported the computer, a mouse, the shimmer device, an additional laptop to complete survey questions on, and a video camera to record the facial expressions of the participant. The study utilized a Dell Precision 7510 laptop, which has an i7 core and a 1 TB 7200 rpm SATA Hard Drive with a 15.6-inch display for users. The camera that was used was a Logitech HD 1080p webcam that was placed on top of the laptop to be near eye level with the study participants. The iMotions software randomized the order of the games that the participants would play. Before playing each game, they watched a short tutorial that explained the features of the game they were about to play and the commands that were used within the game. After watching the tutorial, the participants would listen to calming music and watch a calming scene of a fish swimming in the ocean for 90 seconds to get them back to a baseline level of emotional calm before the start of gameplay. Then they would be dropped into the middle of gameplay to play the game for 10 minutes. After 10 minutes, the participants were administered the GEQ (Game Engagement Questionnaire) to get subjective data about the game they just played (22). The GEQ is a validated questionnaire that utilizes a 3-Likert scale throughout 19 questions. The tutorial, calming video, 10 minutes of gameplay, and questionnaire was repeated for each of the games selected for the study.

After finishing all the tests, participants completed one last survey that had them rank the three games in order of preference and then indicate whether they would purchase this game for gameplay in the future. The participants were also asked to provide a price they would be willing to pay for the game that they just played, ranging from $0 to $50. Participants were then compensated $15 in cash for participating in the experiment and escorted out of the study environment. During the experiment, only the researchers were present in the room in which the study was taking place. The participants did not interact with other participants during the times that they were playing the game for the study.

The iMotions software produced a large variety of potential data measures, such as “eyebrow raise” and “lip curl”. All these values were compared against the baseline data that was collected from the participant to establish a threshold level of significance. The threshold was set to 50%; that is, only measures that showed 50% change were used in the analysis. The values that met the threshold were then summed for the experiment and normalized for the length of each experiment. The final values produced were a percentage of the total values for the experiment that exceeded a 50% threshold from the baseline (greater or lower) for the participant in the experiment (20).

The GEQ was scored on a Likert scale and was added to the final data pool, along with the demographic answers of the participant and their final scoring of the three games based upon their preference. Each data set that was collected was identified by its participant ID to ensure that the proper data was associated with the correct participant.

Due to noise in the data from some of the participants during the biometric measures and incomplete survey data, from the original 40 participants, we only considered those with complete records, which left us with 28 participants for the GEQ and 25 for the biometric dataset. No imputations were performed on these datasets.

The variables from the survey dataset were coded as interval scale variables and the biometric data primarily consisted of continuous variables. The final survey data analyzed was composed of 28 participants’ answers to 19 questions from the GEQ as well as our participants’ preference for the games they played. The dataset was normalized before the analysis of the biometric and subjective measures. For the survey, we looked at the four dimensions, absorption, flow, presence, and immersion, belonging to the GEQ questionnaire (21) and took the mean of the questions as the respective value for that particular dimension. For the analysis, the biometric and survey data identified during and after gameplay were explored to find if there were correlations amongst any of the variables, using Pearson’s correlation coefficients (Table 1).

**Table 1.** Pearson’s correlation coefficient between GEQ dimensions and physiological measures.

| **Physiological Measures** | **Absorption** | **Flow** | **Presence** | **Immersion** |
| --- | --- | --- | --- | --- |
| Anger | 0.05 | 0.08 | -0.09 | 0.08 |
| Sadness | -0.19 | -0.16 | 0.13 | -0.25 |
| Disgust | -0.09 | 0.02 | -0.14 | 0.04 |
| Joy | -0.19 | -0.28 | -0.05 | -0.02 |
| Surprise | -0.05 | -0.01 | -0.08 | 0.08 |
| Fear | 0.20 | 0.23 | -0.02 | 0.17 |
| Contempt | -0.15 | -0.24 | 0.04 | -0.19 |
| Engagement | -0.16 | -0.20 | -0.09 | -0.05 |
| Attention | 0.16 | -0.02 | 0.09 | 0.23 |
| Positive Count | -0.18 | -0.28 | -0.04 | -0.02 |
| Negative Count | -0.15 | -0.10 | -0.02 | -0.18 |
| Neutral Count | 0.24 | 0.10 | 0.11 | 0.24 |
| Brow Furrow | -0.08 | -0.13 | -0.06 | -0.18 |
| Brow Raise | 0.08 | 0.04 | -0.19 | 0.03 |
| Lip Corner Depressor | 0.02 | 0.00 | -0.18 | 0.05 |
| Smile Count | -0.19 | -0.28 | -0.04 | -0.02 |
| Inner Brow Raise | -0.19 | -0.18 | 0.12 | -0.28 |
| Eye Closure Count | -0.22 | -0.31 | -0.37 | -0.11 |
| Nose Wrinkle Count | -0.06 | 0.07 | -0.10 | 0.11 |
| Upper Lip Raise Count | -0.22 | -0.10 | -0.02 | -0.02 |
| Lip Suck Count | -0.10 | -0.10 | -0.07 | 0.06 |
| Lip Press Count | -0.19 | -0.03 | -0.03 | 0.05 |
| Mouth Open Count | 0.03 | 0.21 | 0.04 | 0.23 |
| Chin Raise Count | 0.12 | 0.05 | -0.09 | -0.07 |
| Smirk Count | -0.15 | -0.25 | 0.04 | -0.19 |
| Lip Pucker Count | -0.27 | -0.09 | -0.12 | -0.04 |
| Cheek Raise Count | -0.18 | -0.19 | -0.22 | -0.03 |
| Dimpler Count | -0.19 | -0.07 | 0.01 | 0.10 |
| Eye Widen Count | 0.33 | 0.25 | 0.02 | 0.32 |
| Lid Tighten Count | -0.12 | -0.06 | 0.07 | -0.10 |
| Lip Stretch Count | -0.25 | -0.21 | -0.15 | 0.02 |
| Jaw Drop Count | 0.03 | 0.29 | 0.10 | 0.25 |

For the analysis of the potential to predict self-reported engagement of the three video games, biometric and survey data identified during and after gameplay were explored as potential predictors of engagement. We performed two studies, one with the outcome variable consisting of the game’s rank order of preference and another with the outcome variable consisting of the level of the game of engagement. For engagement, DOOM was coded as 1 (highest engagement), with BattleBorn as 2, and Dead Effect 2 as 3. Preference was ranked as 1, 2, or 3 depending on how the participant ranked the games, with 1 being the most preferred and 3 consisting of the least preferred game. The final survey data analyzed was composed of 28 participants’ answers to 19 questions from the GEQ as well as our participants’ preference for the games they played. The biometric and survey data features considered here are listed in Table 2 and Table 3. Due to the small sample size, as well as our ordinal datasets, only three non-parametric machine learning methods were chosen for the classification of the data, and the results of these three models were compared. The three models considered here are Support Vector Machine (SVM), Bayesian Network, and Random Forests (23).

**Table 2.** Variance Inflation Factors of Survey Dataset for Preference

| **Variable** | **VIF** |
| --- | --- |
| Gamer Experience | 1.675803 |
| Worth | 1.760959 |
| Purchase | 2.410443 |
| *I lose track of time* | 2.177809 |
| *Things seem to happen automatically* | 1.561087 |
| *I feel different* | 3.312574 |
| *I feel scared* | 2.670017 |
| *The game feels real* | 2.612103 |
| *If someone talks to me, I don't hear them* | 4.296267 |
| *I get wound up* | 2.228233 |
| *Time seems to kind of stand still or stop* | 2.467846 |
| *I feel spaced out* | 2.050168 |
| *I don't answer when someone talks to me* | 2.847635 |
| *I can't tell that I'm getting tired* | 1.501123 |
| *Playing seems automatic* | 2.416228 |
| *My thoughts go fast* | 1.630773 |
| *I lose track of where I am* | 2.266367 |
| *I play without thinking about how to play* | 1.914802 |
| *Playing makes me feel calm* | 2.041668 |
| *I play longer than I meant to* | 1.862299 |
| *I really get into the game* | 2.857968 |
| *I feel like I just can't stop playing* | 3.901467 |

**Table 3.** Variance Inflation Factors of Biometric Dataset for Preference

| **Variable** | **VIF** |
| --- | --- |
| *HasPeak* | *1.244416* |
| *PeakCount* | *1.267829* |
| *Stimulus... Marker.Duration...ms* | *1.068957* |
| *Count.Frames.In.Stimulus...Marker* | *1.422789* |
| *Anger.Time.Percent* | *1.238757* |
| *Sadness.Count.Frames...Threshold* | *1.721020* |
| *Disgust.Count.Frames....Threshold* | *1.395853* |
| *Joy.Time.Percent* | *1.083814* |
| *Surprise.Count.Frames...Threshold* | *1.320422* |
| *Fear.Time.Percent* | *1.064214* |
| *Contempt.Time.Percent* | *1.076531* |
| *Attention.Count.Frames...Threshold* | *1.364135* |
| *Negative.Time.Percent* | *2.278113* |

We performed chi-squared tests on our variables to find the corresponding p-values of each variable. Kruskal-Wallis tests were done on our biometric dataset due to our small sample size and our outcome variable consisting of three outcomes in ranked order. We used these p-values of the different predictor variables, the correlation coefficients, and the multicollinearity between the variables to remove variables that were not considered informative. We used the receiver operating characteristic (ROC) measure, the F-Measure, and the accuracy of our models to evaluate their efficacy. Each dataset was divided into 80% training and 20% testing dataset and a ten-cross fold validation were used when implementing the machine learning models.

After the preliminary analysis, we looked at the multicollinearity of the variables to ensure that our results would not be skewed by a variable having a large effect on another one during predictive analysis. Both the survey dataset and biometric datasets were treated as different datasets since they were both being analyzed separately. Our survey dataset did not exhibit much multicollinearity (Table 2), with the highest variance inflation factor (vif=4.30) being for question F: “If someone talks to me I don’t hear them” at 4.30. We did not remove this variable when conducting further analysis tests since our cutoff point for our vif was 5.0 (23).

In Table 3 we present the resulting variables from our biometric dataset after multicollinearity was measured and variables showing large correlation values were removed. It is possible that the variables removed may have had a high correlation due to similar measurements being taken to assess a specific emotion. We were left with 13 variables of the original 29 variables in the biometric dataset after removing those with a vif greater than 5.0 (23). The analysis of the data was performed using R (https://www.r-project.org/) and the machine learning models and predictions were conducted using Weka (24).

## 3. Results

Table 1 shows the correlation coefficients between the GEQ dimensions and the physiological measures. The results depict different patterns and strengths in the covariance. For example, ‘flow’ seems to have the highest covariances with measures from the facial expression analysis, with ‘joy’, ‘contempt’, ‘smile count’, ‘eye closure count’, and ‘smirk count’ having negative R-values and ‘eye widen count’ and ‘jaw drop count’ resulting in positive R-values. Meanwhile, presence was only correlated with one physiological measure, eye closure count (−0.37).

Table 4, depicts the performance for our predictive models for preference using both survey and biometric data. The results show that the biometric dataset performed poorly, with the survey dataset performing well when using two of the three predictive models. From Table 5, it is apparent that the biometric dataset performed better overall than the survey dataset when measuring engagement based on a game’s metrics. Overall, the survey dataset did not do very well in any of our models, with a predictive power not much better than random for any of the given methods. The biometric dataset did outperform the survey dataset in all three of the models. The best performing model can predict engagement with an accuracy of 74.67%, a ROC of 0.876, and an F-Measure of 0.746 using Random Forests. Our Bayesian Network performed slightly worse than that, with an accuracy of 64.67%, a ROC of 0.813 and an F-Measure of 0.651. Our SVM model performed with only a 40.67% accuracy, a ROC of 0.561 and an F-Measure of 0.381.

**Table 4.** Performance of Predictive Models for Preference

| **SURVEY DATASET** |  |  |  |
| --- | --- | --- | --- |
| *Method* | *ROC* | *F-Measure* | *Test Accuracy* |
| *SVM* | *0.879* | *0.802* | *80%* |
| *Bayesian Networks* | *0.707* | *0.504* | *51.33%* |
| *Random Forests* | *0.976* | *0.920* | *92%* |
| **BIOMETRIC DATASET** |  |  |  |
| *Method* | *ROC* | *F-Measure* | *Test Accuracy* |
| *SVM* | *0.472* | *0.317* | *31.33%* |
| *Bayesian Networks* | *0.522* | *0.307* | *33.33%* |
| *Random Forests* | *0.601* | *0.446* | *44.67%* |

**Table 5.** Performance of Predictive Models for Games

| **SURVEY DATASET** |  |  |  |
| --- | --- | --- | --- |
| *Method* | *ROC* | *F-Measure* | *Test Accuracy* |
| *SVM* | *0.563* | *0.388* | *38.095%* |
| *Bayesian Networks* | *0.606* | *0.428* | *42.857%* |
| *Random Forests* | *0.618* | *0.438* | *44.048%* |
| **BIOMETRIC DATASET** |  |  |  |
| *Method* | *ROC* | *F-Measure* | *Test Accuracy* |
| *SVM* | *0.561* | *0.381* | *40.67%* |
| *Bayesian Networks* | *0.813* | *0.651* | *64.67%* |
| *Random Forests* | *0.876* | *0.746* | *74.67%* |

As previously stated, the biometric dataset outperformed our survey dataset and the Random Forest model outperformed the Bayesian Networks and Support Vector Machine models for the biometric dataset when predicting engagement. It is possible that having used a bagging method, such as Random Forests, helped it achieve higher accuracy for the prediction of the success of a game (Kuhn, 2013). When predicting for self-reported preference, however, the survey dataset predicted with much higher accuracy than the biometric dataset.

## 4. Discussion and Conclusions

This study aimed to identify whether there are any correlations between our biometric results and survey results to find if biometric feedback correlates to what an individual reports on a user-reported survey and if the engagement or preference of video games could be predicted using biometric data during gameplay rather than the usual survey data when using machine learning models of success prediction post gameplay. Whereas studies are starting to surface using biometric data during gameplay for video game engagement, there has yet to be a study that compared the usual user testing methods to biometric data to measure the engagement of a participant. This study provides an impetus for much larger and comprehensive studies.

We first analyzed the correlation between our biometric results and our survey results to see if biometric feedback correlates to what an individual reports on a user-reported survey and whether there were any significant differences between each game and the GEQ reported answers from participants. Although there were some negative and positive correlations with our different dimensions of engagement, additional work is needed to establish if comparing user feedback with biometric measures is both feasible as well as reasonable in a ‘game development context’. While no statistical significances were observed between the games and the GEQ measures, having a larger cohort would help in gathering a more accurate measure between the games and the GEQ dimensions.

In this study, we also explored how well we could predict the preference and level of engagement based on metrics of a video game utilizing biometric data, and whether there was an improvement in the prediction of these outcomes when compared to survey data. We collected both survey data and biometric data from a pool of participants both during and after gameplay, and then used our models to see how well we could correctly classify the success of the three video games we studied.

Our results indicated that, overall, biometric data did a better job at indicating the level of engagement of a video game, with a Random Forests prediction model performing as the best for both datasets with an accuracy of 74.67% for the biometric data, whereas survey data managed to achieve an accuracy of 44.048%. These results indicate that biometric data is a better measure of engagement and further analysis of this method of tracking feedback during gameplay should continue to be studied, especially if companies want to create games that will engage the consumer and keep the consumer playing the game for a longer period.

Our results also indicated that for self-reported preference, biometric data was a poor predictor. Self-reported survey data was a better predictor of preference, which would make sense seeing as an individual may report higher levels of engagement to the GEQ questionnaire for a game they prefer. Further analysis, however, is needed to see why the biometric data results were low, as well as why there is a large number of incorrectly predicted preferences when utilizing the biometric dataset to train our model.

One of the main limitations of this study included the small sample size of our data, which was further reduced due to noisy data during gameplay or missing data from participants. Being able to include more participants in our study and in the data being analyzed would allow us to produce more robust results. Another limitation is the possibility of a much more complex and non-linear relationship between the physiological measures and the GEQ dimensions that cannot be captured through typical linear methods.

Further analysis with different algorithms to find the best fit model, as well as understanding the impact of different predictor variables on our model is still needed. Identifying which variables are likely to give us a good estimate, as well as creating a reproducible model would aid in streamlining the testing of games to assess the future success of new and upcoming games. Further analysis of the differences between the levels of game experience in participants should also be done. As noted, consumers that are experienced gamers may react differently than those who have been less exposed to video games and produce different results in the biometric dataset, such as decreased emotional and engagement responses to the video games that they are playing (Mandryk et al., 2006). Additional analysis with other user feedback reports, such as the multi-dimensional user experience survey, iGEQ, may also be worth performing since the GEQ is a one-dimensional survey (Poels et al., 2013).

## Acknowledgments

JCF and JV were partially supported by the National Center for Advancing Translational Sciences of the National Institutes of Health under Award Number UL1TR002538 and the National Library of Medicine T15 LM00712418. Computer resources were provided by the University of Utah Center for High-Performance Computing, which has been partially funded by the NIH Shared Instrumentation grant no. 1S10OD02164401A11. The content is solely the responsibility of the authors and does not necessarily represent the official views of the National Institutes of Health.

## Supplementary Material

### Description of Games Used in this Study

A. Doom DOOM was released on 5/12/2016. It is set in a futuristic universe in a colony on the planet Mars, where the lead character is trying to battle various demons. The game has 1,900,000 downloads on Steam and of those who downloaded it, 8.87% played it during the period of 5/23/17 to 6/6/17. The player reviews were mostly positive for this game.
B. BattleBorn BattleBorn was released on 5/2/2016 on Steam. It is set in a space fantasy setting and is a first-person shooter where you play as a hero character. The game has 321, 642 downloads on Steam and 2.36% of those played it during the period of 5/23/17 to 6/6/17. The player reviews for this game are largely mixed.
C. Dead Effect 2 The last game selected was Dead Effect 2. It was released on 5/6/2016 and it had 60,798 downloads on Steam of which 4.84% of owners played it during the period from 5/23/17 to 6/6/2017. Dead Effect 2 is a futuristic first-person shooter that involves the main character battling enemy soldiers and zombies on a space station. The reviews for the game have been mostly positive.

